# Human LAMP1 accelerates Lassa virus fusion and potently promotes fusion pore dilation upon forcing viral fusion with non-endosomal membrane

**DOI:** 10.1101/2022.06.01.494281

**Authors:** You Zhang, Juan Carlos de la Torre, Gregory B. Melikyan

## Abstract

Lassa virus (LASV) cell entry is mediated by the interaction of the virus glycoprotein complex (GPC) with alpha-dystroglycan at the cell surface followed by binding to LAMP1 in late endosomes. However, LAMP1 is not absolutely required for LASV fusion, as this virus can infect LAMP1-deficient cells. Here, we used LASV GPC pseudoviruses, LASV virus-like particles and recombinant lymphocytic choriomeningitis virus expressing LASV GPC to investigate the role of human LAMP1 (hLAMP1) in LASV fusion with human and avian cells expressing a LAMP1 ortholog that does not support LASV entry. We employed a combination of single virus imaging and virus population-based fusion and infectivity assays to dissect the hLAMP1 requirement for initiation and completion of LASV fusion that culminates in the release of viral ribonucleoprotein into the cytoplasm. Unexpectedly, ectopic expression of hLAMP1 accelerated the kinetics of small fusion pore formation, but only modestly increased productive LASV fusion and infection of human and avian cells. To assess the effects of hLAMP1 in the absence of requisite endosomal host factors, we forced LASV fusion with the plasma membrane by applying low pH. Unlike the conventional LASV entry pathway, ectopic hLAMP1 expression dramatically promoted the initial and full dilation of pores formed through forced fusion at the plasma membrane. We further show that, while the soluble hLAMP1 ectodomain accelerates the kinetics of nascent pore formation, it fails to promote efficient pore dilation, suggesting the hLAMP1 transmembrane domain is involved in this late stage of LASV fusion. These findings reveal a previously unappreciated role of hLAMP1 in promoting dilation of LASV fusion pores, which is difficult to ascertain for endosomal fusion where several co-factors, such as bis(monoacylglycero)phosphate, likely regulate LASV entry.

**Author Summary:** Lassa virus (LASV) enters cells *via* fusion with acidic endosomes mediated by the viral glycoprotein complex (GPC) interaction with the intracellular receptor LAMP1. However, the requirement for LAMP1 is not absolute, as LASV can infect avian cells expressing a LAMP1 ortholog that does not interact with GPC. To delineate the role of LAMP1 in LASV entry, we developed assays to monitor the formation of nascent fusion pores, as well as their initial and complete dilation to sizes that allow productive infection of avian cells by LASV GPC pseudoviruses. This novel approach provided unprecedented details regarding the dynamics of LASV fusion pores and revealed that ectopic expression of human LAMP1 in avian cells leads to a marked acceleration of fusion but modestly increases the likelihood of complete pore dilation and infection. In contrast, human LAMP1 expression dramatically enhanced the propensity of nascent pores to fully enlarge when LASV fusion with the plasma membrane was forced by exposure to low pH. We conclude that, whereas the role of LAMP1 in LASV fusion is confounded by an interplay between multiple endosomal factors, the plasma membrane is a suitable target for mechanistic dissection of the roles of endosomal factors in LASV entry.

## Introduction

LASV is an Old World mammarenavirus that infects a broad host range of cells from different species. LASV cell entry is mediated by the viral surface glycoprotein complex (GPC), a trimeric class I fusion protein that consists of non-covalently associated surface (GP1) and transmembrane (GP2) subunits (reviewed in [1, 2]). The GP1 and GP2 subunits are generated through cleavage of the GPC precursor by the host cell protease subtilisin kexin isozyme-1(SKI-1)/site 1 protease (S1P). GP1 is involved in receptor binding and GP2 in membrane fusion. A unique feature of arenavirus GPC proteins is that their stable signal peptide (SSP), which is cleaved off the GP precursor, remains associated with the GPC and plays an important regulatory role in low pH-induced conformational changes in GPC that lead to fusion of the viral and cell membranes [3–8]. LASV GPC attachment to the alpha-dystroglycan receptor on the cell surface leads to virus internalization and transport to acidic endosomes where low pH promotes virus dissociation from alpha-dystroglycan and attachment to the intracellular receptor, LAMP1. LAMP1, a marker for late endosomes/lysosomes, has been shown to serve as a specific receptor for LASV and not for other mammarenaviruses, such as lymphocytic choriomeningitis virus (LCMV) [9]. Human LAMP1 (hLAMP1), but not human LAMP2 or avian LAMP1, promotes LASV fusion with late endosomes [9].

Recent studies by others and our group have shown that hLAMP1, while promoting LASV fusion and infection, is not absolutely required for virus entry, since cells lacking human LAMP1 support basal levels of LASV fusion/infection [10–12]. The ability of LASV GPC to mediate membrane fusion in the absence of hLAMP1 is consistent with the reports that sufficiently acidic pH induces irreversible GPC conformational changes leading to shedding of the GP1 subunit [9, 12, 13]. Mechanistic studies revealed that hLAMP1 binding shifts the pH-optimum for GPC-mediated fusion to a higher pH [10, 12]. Thus, LASV fusion with hLAMP1 expressing cells is likely initiated in less acidic maturing endosomal compartments, prior to virus delivery into late endosomes/lysosomes.

Our recent study identified a novel co-factor required for completion of LASV fusion – the late-endosome-resident lipid, bis(monoacylglycero)phosphate (BMP) [12]. BMP specifically and potently promotes the late stages of LASV GPC-mediated fusion – formation and enlargement of fusion pores. Whereas hLAMP1 binding clearly augments low pH-dependent refolding of LASV GPC, whether this intracellular receptor also modulates late steps of virus fusion remains unclear. Using a cell-cell fusion model, we have shown that hLAMP1 overexpression facilitates transition from hemifusion (merger of contacting membrane leaflets without fusion pore formation) to full fusion [12]. However, the role of hLAMP1 in controlling distinct steps of virus-endosome fusion has not been elucidated.

Here, we employed a combination of single LASV pseudovirus (LASVpp) tracking in live cells alongside bulk virus-cell fusion and infectivity assays to assess the effect of hLAMP1 on distinct steps of viral fusion. Pseudoviruses co-labeled with viral content marker and an internal pH sensor enable detection of single virus fusion events and monitoring the initial enlargement of fusion pores, whereas bulk fusion/infectivity assay report functional enlargement of fusion pores. We find that hLAMP1 overexpression in both human and avian cell lines moderately promotes fusion and infection through an endocytic entry pathway, whereas LASV GPC-mediated fusion at the cell surface forced by exposure to low pH was dramatically enhanced and accelerated in hLAMP1-expressing cells. Real-time imaging of forced LASVpp fusion with avian cells revealed a strong enhancement in fusion pore dilation by upon hLAMP1 expression. Our results thus provide new insights into the role of hLAMP1 in early and late stages of LASV fusion and reveal key differences in permissiveness of endosomes and the plasma membrane for LASV fusion.

## Results

### Ectopic expression of hLAMP1 potentiates LASV fusion and infection of human and avian cells

We used two cell lines to examine the role of LAMP1 in LASV fusion – human lung epithelial A549 cells endogenously expressing human LAMP1 (hLAMP1) and chicken DF-1 fibroblasts expressing the avian LAMP1 ortholog which does not support LASV entry [9]. Immunofluorescence staining confirmed that A549 cells expressed a basal level of hLAMP1, in agreement with our immunoblotting data [12], whereas this protein was not detectable in DF-1 cells (Fig. 1). We next stably transduced A549 and DF-1 cells with wild-type hLAMP1 (LAMP1-WT) or with the LAMP1-d384 mutant which lacks an endocytic signal and is primarily expressed on the cell surface [9]. Transduction with hLAMP1 yielded robust levels of expression in both A549 and DF-1 cells, with hLAMP1 primarily distributed to intracellular compartments (Fig. 1A-D). To assess the levels of hLAMP1 on the cell surface, immunofluorescence staining was performed without cell membrane permeabilization. This protocol revealed modest levels of LAMP1-WT on the surface of both A549 and DF-1 cell lines overexpressing LAMP1-WT, whereas the surface levels of LAMP1-d384 were much higher, reaching approximately half of the total expressed hLAMP1 (Fig. 1A-D).

**Figure 1.**
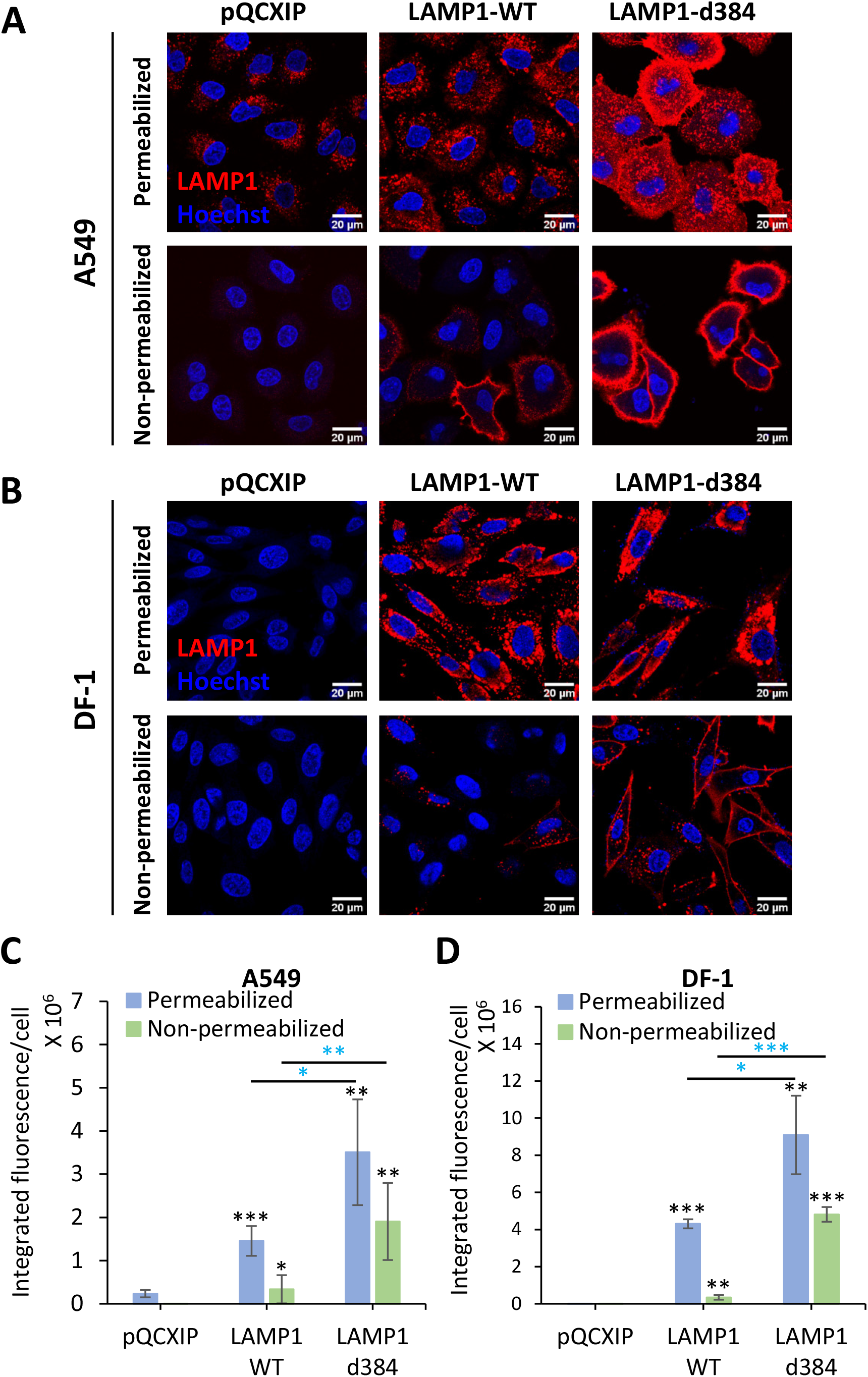
Analysis of endogenous and ectopic expression and localization of hLAMP1 in A549 and DF-1 cells. A549 or DF-1 cells were transduced with an empty pQCXIP vector (control) or vector expressing human LAMP1-WT or the LAMP1-d384 mutant. Cells were fixed and whether left untreated (bottom panels in A and B) or permeabilized with 125 µg/ml digitonin and immunostained for human LAMP1. **(A, C)** Images and quantification of hLAMP1 expression in A549 cells. Images acquired under same exposure conditions, but the brightness and contrast settings in panel A and B are different to ensure optimal display. **(B, D)** Images and quantification of hLAMP1 expression with DF-1 cells. Data shown are means ± SD of five fields of view for each condition. Blue asterisks show significance levels for the difference between LAMP1-WT and LAMP1-d384. Black asterisks on the top of bars represent significance relative to the vector control. Data were analyzed by Student’s t-test. *, p<0.05; **, p<0.01; ***, p<0.001.

Consistent with the requirement for hLAMP1, infection of DF-1 cells by HIV-1 particles pseudotyped with LASV GPC (LASVpp) measured by a single-cycle infectivity assay was ∼100 fold lower than infection of A549 cells (Fig. 2A). To rule out the presence of species-specific endosomal restriction factors in DF-1 cells, we bypassed endosomal entry of LASVpp by triggering virus fusion with the plasma membrane through low pH exposure (referred to as forced fusion, see Materials and Methods for details). Prior to adding the virus and triggering fusion by acidic pH, cells were pre-treated with Bafilomycin A1 (BafA1) to raise endosomal pH and block the endosomal entry of LASVpp. The forced fusion protocol resulted in ∼10-fold less efficient LASVpp infection in both DF-1 and A549 cells compared to endosomal entry (Fig. 2A). Thus, DF-1 cells are poorly permissive for LASVpp infection and forced LASV GPC-mediated fusion with the plasma membrane does not enable efficient infection of human or avian cells compared to a conventional entry route.

**Figure 2.**
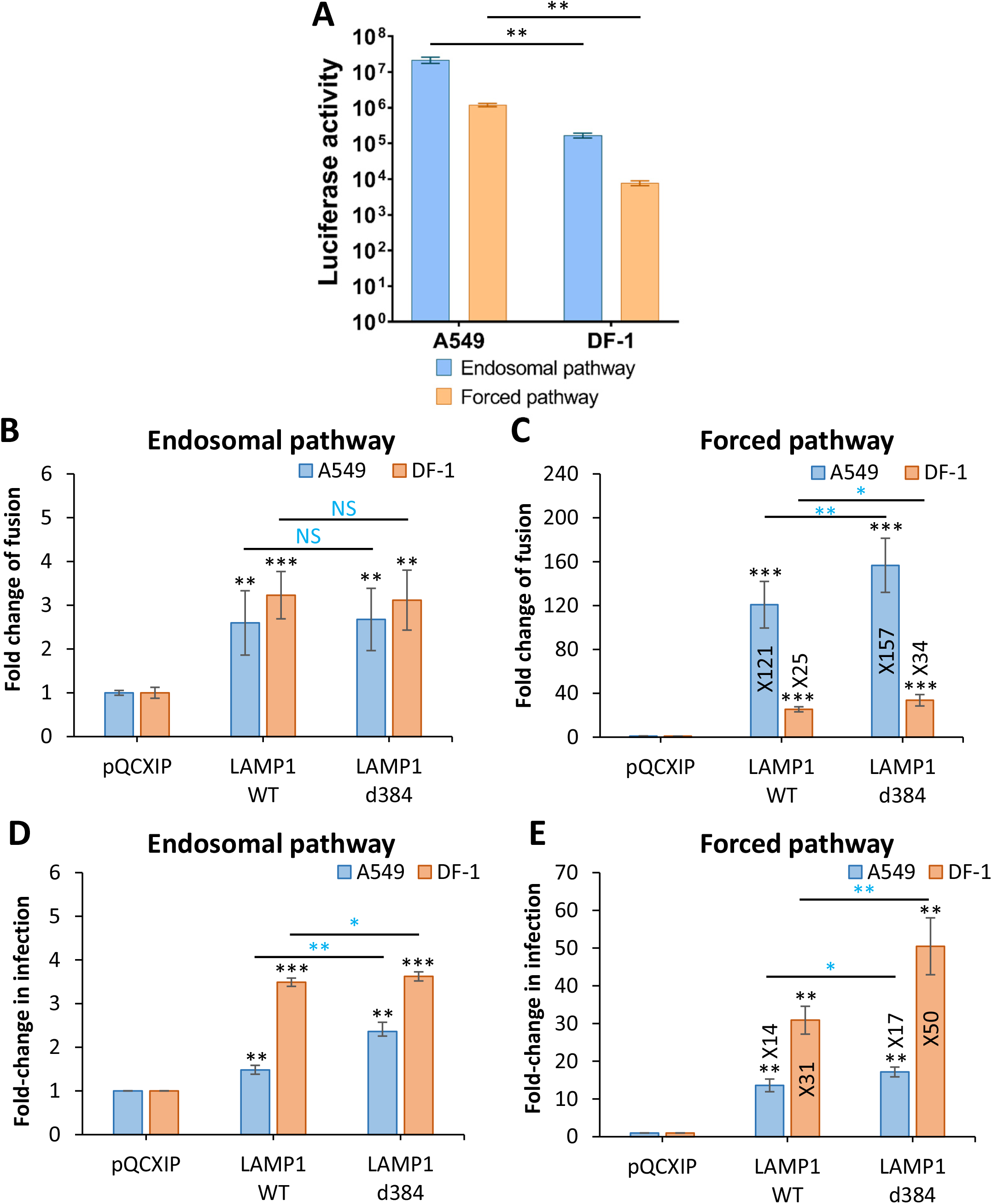
LAMP1 expression enhances LASVpp fusion and infection in A549 and DF-1 cells. **(A)** Efficiency of A549 and DF-1 cell infection through endosomal and forced fusion protocols. **(B)** LASVpp-BlaM fusion with A549 and DF-1 cells. LASVpp entry through an endosomal pathway was initiated by pre-binding pseudoviruses in the cold, shifting to 37°C and incubating for 2 h. **(C)** Low pH-forced fusion of LASVpp with A549 and DF-1 cells. Cells were pretreated with 0.2 µM BafA1 for 1 h prior to binding pseudoviruses in the cold. Fusion was triggered by applying pH 5.0 citrate buffer at 37 °C for 20 min followed by additional incubation in a neutral pH medium at 37 °C for 30 min. **(D)** LASVpp infection of A549 and DF-1 cells. Luciferase-encoding LASVpp were bound to cells in cold. Cells were incubated in 37 °C for 36 h to allow infection. **(E)** LASVpp infection through low pH bypass protocol in A549 and DF-1 cells described in panel C. After forced fusion, cells at 37 °C for 36 h before reading the resulting luciferase signal. Data shown are means ± SD of three independent experiments. Data were analyzed by Student’s t-test. *, p<0.05; **, p<0.01; ***, p<0.001; NS, not significant. Blue asterisks show the significance levels for the difference between LAMP1-WT and LAMP1-d384, black asterisks on the top of bars represent significance relative to the vector control.

Next, we examined the effect of ectopic hLAMP1 expression on the efficiency of LASVpp fusion with A549 and DF-1 cells using a direct virus-cell fusion assay that reports the delivery of virus-incorporated beta-lactamase (BlaM) into the cytoplasm [14, 15]. Pseudoviruses containing BlaM-Vpr chimera (Fig. S1A) were bound to DF-1 or A549 cells and the efficiency of viral fusion was assessed by the resulting cytoplasmic BlaM activity. Consistent with the infectivity data (Fig. 2A), the LASVpp fusion signal was ∼10-fold lower in DF-1 cells lacking hLAMP1 compared to A549 cells (data not shown). Ectopic expression of LAMP1-WT or LAMP1-d384 resulted in a modest, approximately 3-fold, increase in virus fusion and single-cycle infection of both A549 and DF-1 cells (Fig. 2B, D), consistent with the notion that LAMP1 is dispensable for LASV entry.

### hLAMP1 expression dramatically promotes forced LASV pseudovirus fusion with the plasma membrane

A modest increase in LASVpp fusion and infection upon hLAMP1 expression indicates that endogenous levels of this receptor in A549 cells may be sufficient for near-optimal LASVpp entry. On the other hand, the lack of a strong effect on viral fusion and infection of DF-1 cells may be indicative of a limited access of LASVpp to the ectopically expressed endosomal hLAMP1. To probe the effects of hLAMP1 on LASVpp fusion with membranes that do not contain significant amounts of this intracellular receptor and to ensure a more tractable system for viral fusion, we bypassed the internalization and endosomal trafficking steps by forcing LASVpp fusion with the plasma membrane through exposure to low pH. The forced LASVpp fusion measured by the BlaM assay was dramatically enhanced in both A549 and DF-1 cells upon hLAMP1 expression (Fig. 2C). Expression of LAMP1-WT or LAMP1-d384 in DF-1 cells caused a somewhat less dramatic (∼30-fold) increase in LASVpp fusion compared to A549 cells in which ∼120-160-fold increase in signal was detected. The more dramatic effect of hLAMP1 expression on forced LASVpp fusion with A549 vs DF-1 cells (Fig. 2C, E) was unexpected, given that the former cells express endogenous hLAMP1. The striking increase in the efficiency of forced LASVpp fusion is in agreement with the marked enhancement of LASV GPC-mediated cell-cell fusion upon ectopic expression of hLAMP1 in DF-1 cells [12]. Consistent with the higher expression of the mutant LAMP1 on cells (Fig. 1), forced fusion with both A549 and DF-1 cells expressing LAMP1-d384 was significantly more efficient than fusion with cells expressing LAMP1-WT (Fig. 2C). Thus, ectopic hLAMP1 expression greatly enhances the otherwise sub-optimal low pH-mediated LASVpp fusion with the plasma membrane, in contrast to fusion with endosomes which is less affected by hLAMP1 overexpression.

The markedly more potent effect of ectopic hLAMP1 expression on the forced LASVpp fusion compared to endosomal entry into DF-1 cells supports the notion that this virus may have limited access to endosomal hLAMP1 in these cells. Similar to the effect of hLAMP1 overexpression on forced viral fusion, surface expression of this receptor also strongly enhanced the forced LASVpp infection of both cell types (Fig. 2E). Interestingly, the relative magnitude of the infection-enhancing effect of hLAMP1 in A549 vs DF-1 cells was reversed compared to the fusion assay (Fig. 2B). Here, a greater gain in infection (up to 50-fold) was observed in DF-1 cells compared to A549 cells (∼15-fold, Fig. 2E). It is also noteworthy that hLAMP1 expression had a commensurate effect on viral fusion and infection in DF-1 cells, whereas an increase in fusion efficiency was much more pronounced compared to infection in A549 cells (Fig. 2C, E). A discordance between fold-enhancement of forced fusion *vs* infection in hLAMP1 expressing A549 cells might be due to a less efficient infection following the GPC-mediated fusion of HIV-1 pseudoviruses with the plasma membrane as compared to fusion with endosomes.

### hLAMP1 enhances fusion and infection of LASV VLPs and recombinant Lassa viruses

In order to validate the results obtained with LASVpp, we measured the fusion of arenavirus virus-like particles (VLPs) bearing LASV GPC (referred to as LASV-VLP). VLPs were made by co-expressing the New World Junin arenavirus NP and Z proteins with LASV GPC, essentially as described in [16]. To measure VLP-cell fusion, we constructed a JUNV NP-beta-lactamase chimera and incorporated it into LASV-VLPs (Fig. S1). Similar to LASVpp fusion (Fig. 2B), LASV-VLP fusion with A549 and DF-1 cells measured by the NP-BlaM delivery into the cytoplasm was modestly augmented by ectopic expression of LAMP1-WT or LAMP1-d384 (Fig. S2A). Also, similar to LASVpp, forced LASV-VLP fusion with DF-1 and A549 cells expressing LAMP1-WT or LAMP1-d384 was dramatically (20-40-fold) enhanced compared to control cells (Fig. S2B). We note that forced LASV-VLP fusion with A549 cells was less markedly increased upon hLAMP1 expression than forced fusion of LASVpp (compare Fig. 2C and Fig. S2B). These results further demonstrate that, surprisingly, the enhancement of GPC mediated fusion by hLAMP1 is more pronounced at the plasma membranes relative to fusion with endosomes.

Finally, we examined the effect of hLAMP1 expression on infection by recombinant LCMV viruses bearing LASV GPC (LCMV-LASV GPC) [17, 18]. A549 and DF-1 cells transduced with LAMP1-WT, LAMP1-d384 or an empty vector were inoculated with LCMV-LASV GPC, and infection was quantified by microscopy after immunostaining for the LCMV NP protein (Fig. 3A). Consistent with our results with LASVpp infection (Fig. 2D), ectopic expression of hLAMP1 modestly increased recombinant LASV infection in both cell lines (Fig. 3B). However, unlike the forced LASVpp and LASV-VLP fusion and infection, which were dramatically enhanced in cells overexpressing hLAMP1 (Fig. 2C, E and Fig. S2B), forced LCMV-LASV GPC infection increased only ∼6-fold in DF-1 and A549 cells (Fig. 3C). A more potent enhancement of forced LCMV-LASV GPC infection relative to conventional infection is consistent with a limited virus access to ectopically expressed hLAMP1 in endosomes. It is worth noting that, while the magnitude of hLAMP1 effects on fusion/infection of pseudovirus, VLP and recombinant arenavirus varies, forced fusion at the plasma membrane is consistently more dependent on ectopic hLAMP1 expression than fusion through a conventional endosomal entry pathway.

**Figure 3.**
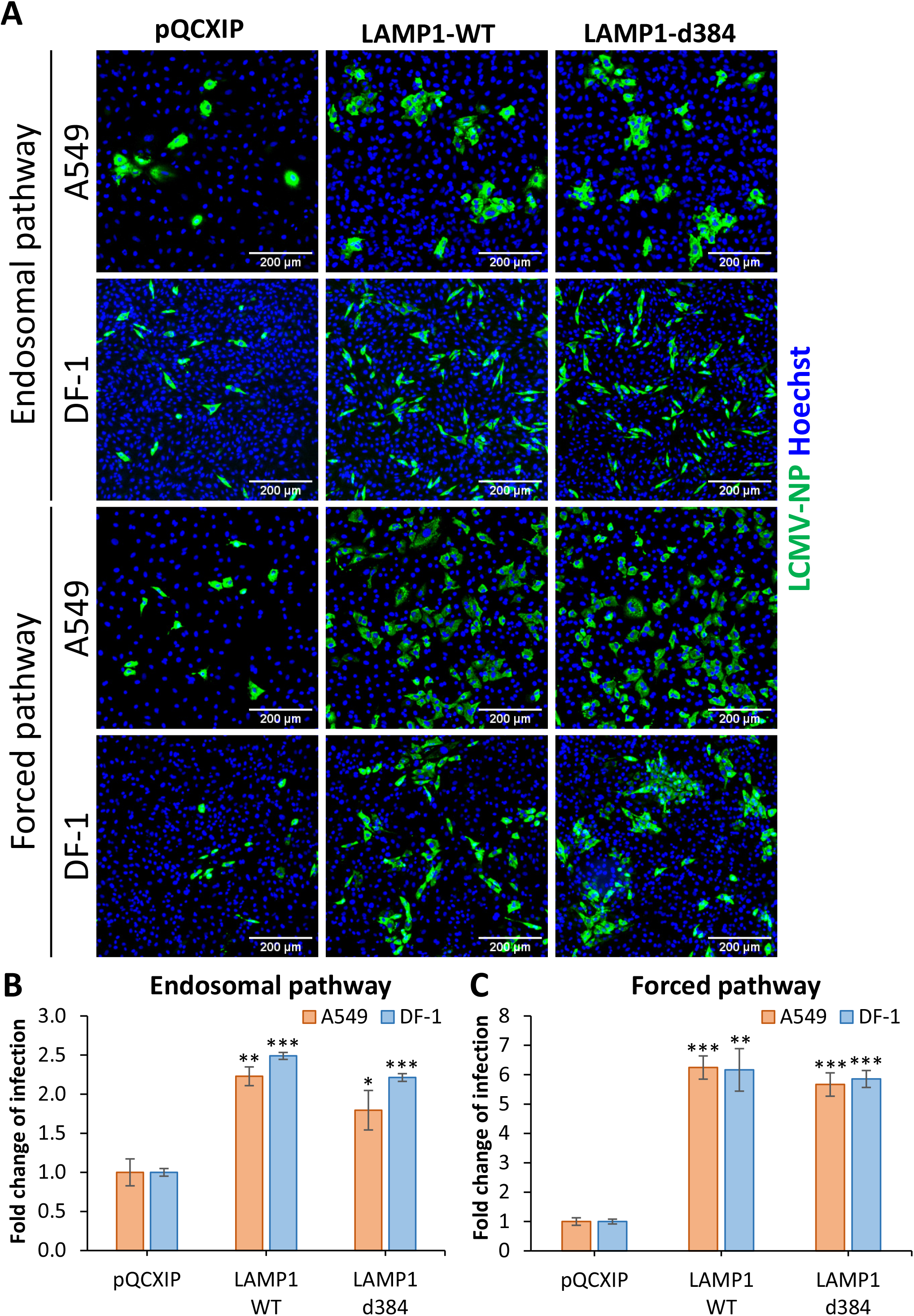
LAMP1 expression enhances recombinant LCMV/LASV-GPC infection of A549 and DF-1 cells. **(A)** Recombinant LCMV/LASV-GPC infection in A549 and DF-1 cells. Cells were allowed to bind LCMV/LASV-GPC in the cold and subsequently incubated at 37 °C for 36 h. For infection in A549 and DF-1 cells achieved through forcing viral fusion by low pH exposure, cells were pretreated with 0.2 µM BafA1 for 1 h prior to binding LCMV/LASV-GPC in the cold. Fusion was triggered by applying pH 5.0 citrate buffer at 37 °C for 20 min followed by additional incubation in a neutral pH medium at 37 °C for 36 h. Infection was detected by immunostaining for LCMV nucleoprotein (NP). **(B)** Quantification of the fold change of the infection through an endosomal pathway. **(C)** Quantification of the fold change of the infection through forced fusion. Data shown are means ± SD of three independent experiments. Data were analyzed by Student’s t-test. *, p<0.05; **, p<0.01; ***, p<0.001.

### Single LASV pseudovirus fusion proceeds through a viral membrane permeabilization step, irrespective of cell type or hLAMP1 expression levels

We next sought to determine which steps of viral fusion are facilitated by hLAMP1 expression by single virus tracking. LASVpp were labeled with the mCherry-2xCL-YFP-Vpr construct, which is incorporated into HIV-1 pseudoviruses and is cleaved by the viral protease upon virus maturation, producing free mCherry and YFP-Vpr [19]. Loss of mCherry signal from the viral particle entering a cell reflects mCherry release into the cytoplasm through a fusion pore, whereas YFP-Vpr is retained in the viral core for a considerable time and thus serves as a reference marker for reliable detection of single fusion events. Also, importantly, the pH-sensitive YFP fluorescence is quenched at low pH [20], thereby reporting changes in intraviral pH. Using this marker, we have previously shown that LASVpp fusion is preceded by a drop in intraviral pH due to permeabilization of the viral membrane in acidic endosomes [20]. This loss of a barrier function of the viral membrane prior to fusion is manifested in YFP fluorescence (green) quenching, whereas subsequent virus fusion results in simultaneous loss of mCherry (red) and recovery of YFP signal due to re-neutralization of the viral interior connected to the cytoplasm through a fusion pore (termed type II fusion, Fig. 4A). Viral membrane permeabilization is likely caused by conformational changes in LASV GPC and requires virus-cell contact prior to exposure to low pH (Zhang and Melikyan, unpublished data). Thus, dual labeling of pseudoviruses with the content marker, mCherry, and the viral core-associated pH sensor, YFP-Vpr, enables sensitive detection of nascent proton-permeable fusion pores and their initial dilation that allows mCherry escape.

**Figure 4.**
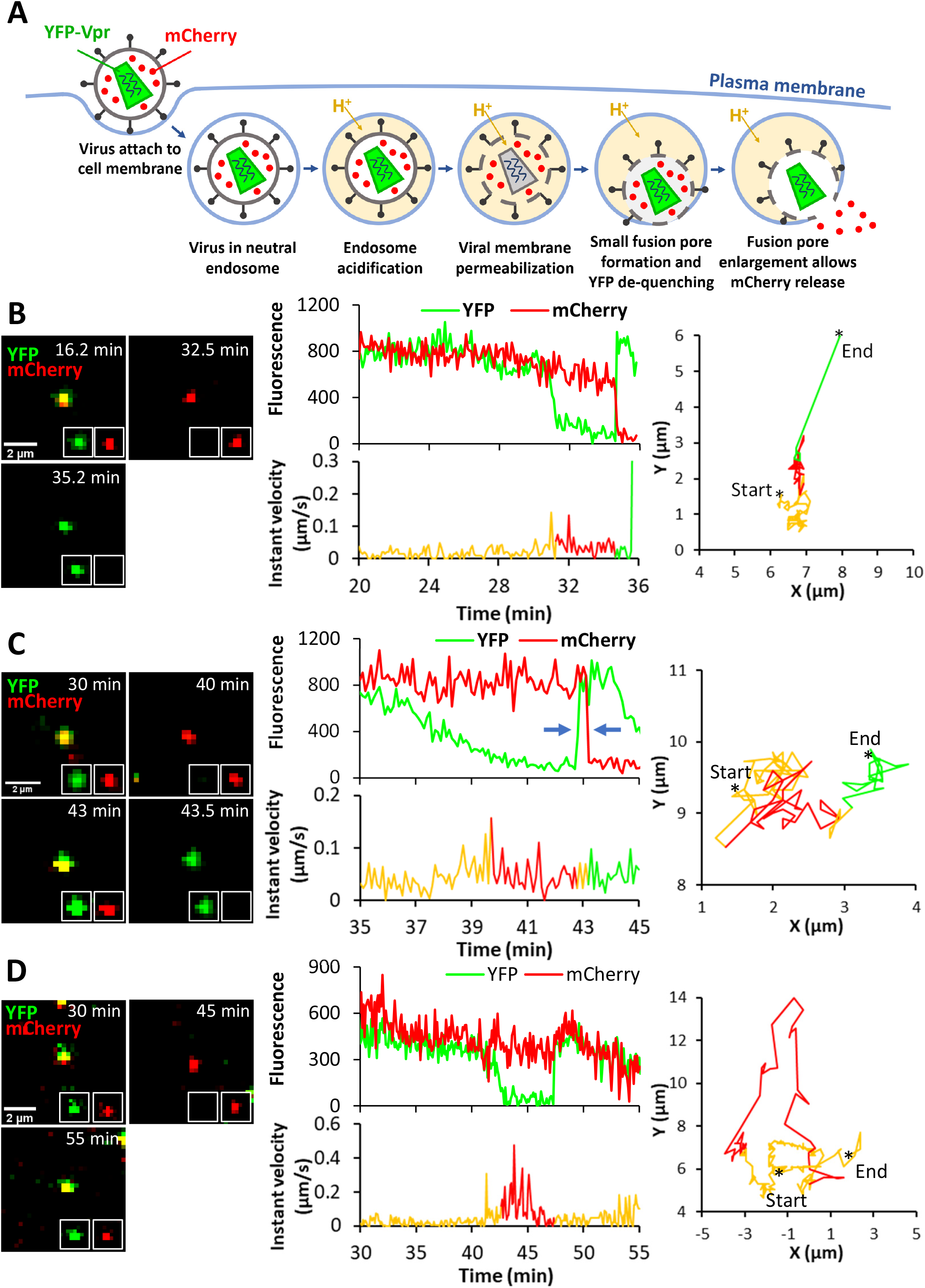
Single LASVpp fusion with DF-1 cells. **(A)** Illustration of fusion of mCherry-2xCL-YFP-Vpr labeled single LASVpp in an acidic endosome. An increase in the virus membrane permeability leads to quenching of the intraviral YFP signal (green) in acidic environment. Subsequent virus fusion with the endosomal membrane results in a loss of mCherry signal (red) through a fusion pore and concomitant re-neutralization of virus interior, as evidenced by YFP signal dequenching. **(B)** LASVpp fusion events (YFP dequenching) with instant mCherry release (quick fusion pore dilation). Time lapse images (left), fluorescence traces (middle top), instant velocity (middle bottom) and trajectory (right) of single LASVpp fusion with DF-1-LAMP1-WT cell showing YFP quenching at 31.3 min and YFP dequenching/mCherry loss at 34.7 min corresponding to virus interior acidification and fusion, respectively (see Movie S1). **(C)** LASVpp fusion events with delayed mCherry release relative to YFP dequenching. Time lapse images (left), fluorescence traces (middle top), instant velocity (middle bottom) and trajectory (right) of single LASVpp fusion with a DF-1-LAMP1-WT cell showing YFP quenching at 39.7 min, YFP dequenching at 42.7 min and mCherry loss at 43.2 min (arrows), indicating virus interior acidification, small fusion pore formation and fusion pore dilation to a diameter exceeding the size of mCherry, respectively (see Movie S2). **(D)** LASVpp fusion events (YFP dequenching) without mCherry release. Time lapse images (left), fluorescence traces (middle top), instant velocity (middle bottom) and trajectory (right) of single LASVpp fusion with a DF-1-LAMP1-WT cell showing YFP quenching at 42.6 min and dequenching at 47.4 min without mCherry loss (see Movie S3).

Using the aforementioned virus labeling strategy, we imaged single LASVpp fusion with DF-1 cells expressing or lacking hLAMP1. DF-1 cells were chosen because of their low permissiveness to LASV fusion/infection in the absence of ectopically expressed hLAMP1 and because, in these cells, the effects of hLAMP1 expression on fusion and infection were more consistent across different virus platforms (LASVpp, LASV-VLP and LCMV-LASV GPC) than in A549 cells (Figs. 2 and 3 and Fig. S2). Similar to our previous finding that LASVpp undergoes type II fusion in A549 cells [20], all single LASVpp fusion events in DF-1 cells exhibited a type II phenotype, regardless of the hLAMP1 expression (Fig. 4B, C and Fig. S3). Interestingly, a fraction of single LASVpp fusion events exhibited a delayed mCherry release relative to YFP dequenching, which marks the opening of a nascent fusion pore (Fig. 4C). This lag in mCherry release ranged from several seconds to minutes (see below) and was most likely caused by delayed enlargement of nascent fusion pores to sizes (∼4 nm) that allowed mCherry release. Thus, YFP dequenching in the context of type II virus-endosome fusion provides a highly sensitive means to detect very small fusion pores, whereas mCherry release is contingent on pore enlargement to a diameter exceeding ∼4 nm.

In addition to type II fusion events culminating in mCherry loss, we also observed YFP quenching and subsequent dequenching without mCherry release for as long as we tracked viral particles (Fig. 4D). The lack of mCherry release could be due to a failure of nascent fusion pores to enlarge and allow mCherry release or due to a full fusion of immature viral particles in which the mCherry marker was not cleaved off the core marker by the HIV-1 protease [19]. Indeed, almost 20% of particles contained uncleaved mCherry-YFP-Vpr construct, as judged by the lack of mCherry release upon saponin lysis *in vitro* (Fig. S4). However, the augmentation of “instantly” dilating pores but not pores that failed to release mCherry upon ectopic hLAMP1 expression (see Fig. 7A below) argues against this possibility. Another reason for YFP dequenching without mCherry release could be virus recycling to the cell surface. However, events that did not culminate in mCherry release were also observed upon low pH-forced virus fusion with the plasma membrane where low pH was maintained throughout the experiment (see below), suggesting that virus cycling to the cell surface is less likely to be responsible for YFP dequenching. We therefore conditionally refer to YFP dequenching without mCherry release as stalled fusion.

Regardless of the timing of mCherry release relative to YFP dequenching, LASVpp fusion was associated with a mixed diffusive and directional motion pattern (Fig. 4), which is typical for endosomal trafficking of internalized cargo and viruses (e.g., [21]).

### hLAMP1 overexpression accelerates single LASV pseudovirus fusion with endosomes without affecting dilation of fusion pores

We observed three single LASVpp fusion phenotypes – instant and delayed pore enlargement and stalled fusion – in DF-1 cells, irrespective of ectopic hLAMP1 expression (Fig. 5A). All three types of single fusion events were promoted ∼4-6-fold upon hLAMP1 expression, in general agreement with the bulk LASVpp fusion results (Fig. 2B). It should be stressed, however, that hLAMP1 expression did not significantly alter the relative weights of the three types of fusion events, including the “instantly” dilating pores (Fig. 5A, *Inset*). Another approach to evaluate the effect of hLAMP1 on the propensity of fusion pores to enlarge is to assess the time required for nascent pore (detected by YFP dequenching) to dilate to sizes that allow mCherry release. “Instant” pore enlargement was defined as simultaneous (within our temporal resolution of 6 sec) YFP dequenching and mCherry loss. The lag time to mCherry release was not significantly affected by hLAMP1 expression (Fig. 5B), in excellent agreement with the finding that hLAMP1 did not significantly increase the fraction of “instant” mCherry release events (Fig. 5A, *Inset*). Thus, ectopic expression of hLAMP1 does not noticeably promote enlargement of fusion pores formed between LASVpp and endosomes.

**Figure 5.**
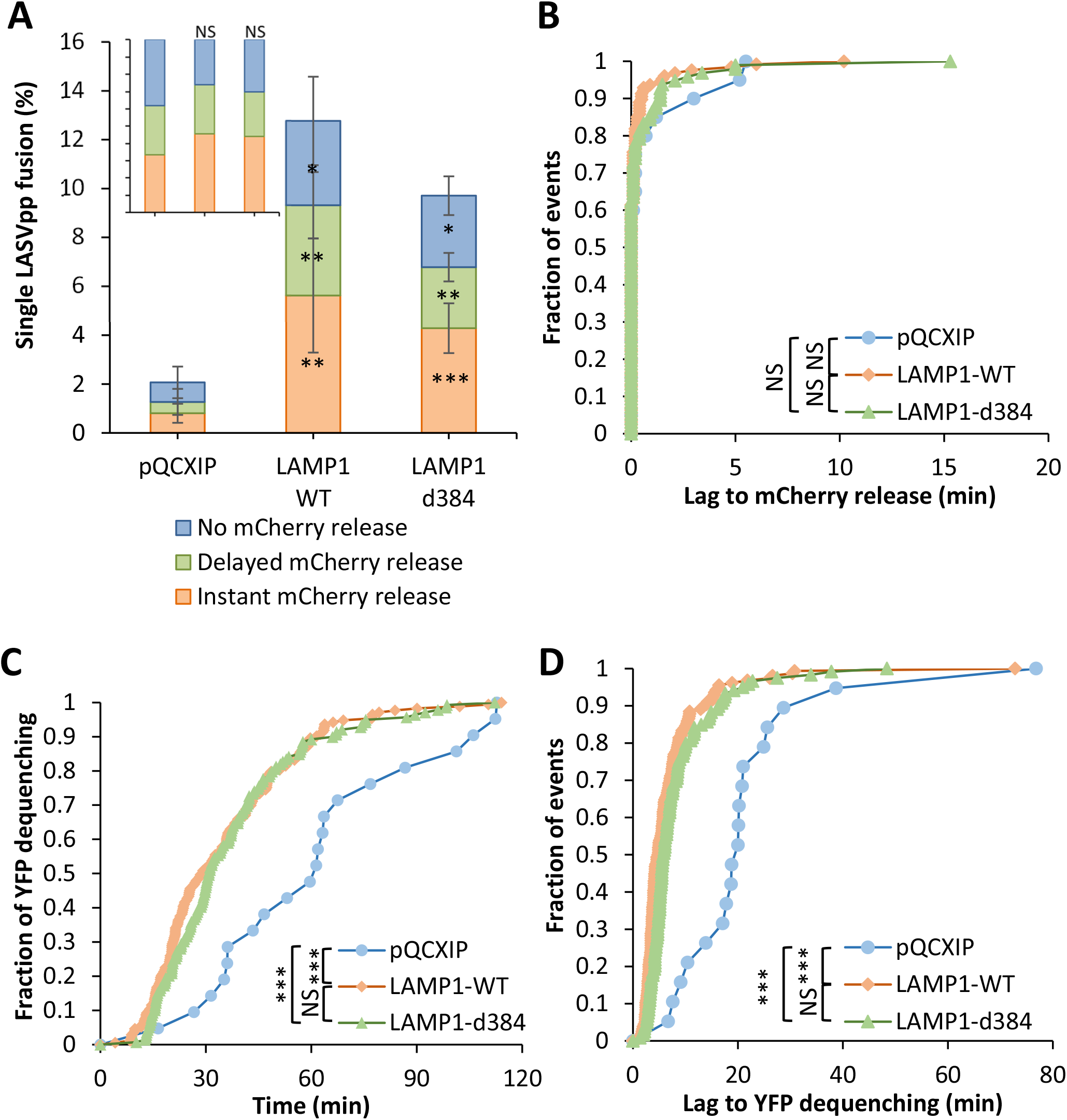
Human LAMP1 expression promotes single LASVpp fusion with DF-1 cells. **(A)** Efficiencies of single LASVpp fusion with instant mCherry release, delayed mCherry release, and without mCherry release in pQCXIP, LAMP1-WT and LAMP1-d384 DF-1 cells. Data shown are means ± SD of 5 independent experiments. Asterisks inside the bars represent significance relative to the vector control. Differences between LAMP1-WT and LAMP1-d384 are not statistically significant for all three categories of fusion. *Inset*: normalized fractions of each category of fusion. **(B)** The distribution of lag times between small fusion pore formation (YFP dequenching) and pore enlargement (loss of mCherry) for LASVpp fusion with DF-1 pQCXIP, LAMP1-WT and LAMP1-d384 cells. **(C)** Kinetics of small pore formation (YFP dequenching) for single LASVpp fusion events in control and hLAMP1 expressing DF-1 cells. **(D)** The distribution of lag times between LASVpp membrane permeabilization (YFP quenching) and small fusion pore formation (YFP dequenching) for LASVpp fusion with DF-1 pQCXIP, LAMP1-WT and LAMP1-d384 cells. Normalized fractions of different single virus fusion events were analyzed by Fisher’s exact test using R Project. Data of lag time between YFP dequenching and mCherry release was analyzed by non-parametric Mann-Whitney test using GraphPad. Other results were analyzed by Student’s t-test. *, p<0.05; **, p<0.01; ***, p<0.001; NS, not significant.

Analysis of the kinetics of nascent fusion pore formation (YFP dequenching) revealed a ∼2-fold increase in the fusion rate with DF-1 cells expressing LAMP1-WT or LAMP1-d384 compared to control cells (Fig. 5C). The faster kinetics of single LASVpp fusion with hLAMP1-expressing DF-1 cells was not caused by more efficient or faster virus endocytosis and delivery into acidic endosomes. Neither the fraction of particles exhibiting YFP quenching (i.e., virus interior acidification in acidic endosomes irrespective of fusion) nor the kinetics of YFP quenching were dependent of hLAMP1 expression (Fig. S5A, B). The fact that only ∼15% of cell-bound particles exhibited YFP quenching in acidic endosomes of DF-1 cells expressing or lacking hLAMP1 is consistent with slow/inefficient endocytosis and virus delivery into late endosomes/lysosomes enriched in LAMP1. Importantly, hLAMP1 expression markedly shortened the half-time of lag between virus interior acidification and small pore formation (YFP dequenching) from ∼18 min to ∼8 min (Fig. 5D). The large lag to small fusion pore formation after virus entry into acidic compartments is thus likely responsible, at least in part, for the slower LASVpp fusion in cells lacking hLAMP1 (Fig. 5C).

### hLAMP1 expression accelerates forced fusion of single LASV pseudoviruses at the cell surface and promotes initial dilation of fusion pores

We have previously shown that hLAMP1 overexpression allows LASV GPC-mediated fusion to occur at higher pH [10, 12]. To control the pH that triggers LASV GPC conformational changes, we employed the forced fusion protocol (Fig. 6A). Fluorescent LASVpp were prebound to DF-1 cells in the cold and their fusion was triggered at 37 °C by applying pH 5.0 buffer. Low pH promoted efficient single LASVpp fusion with control and hLAMP1 expressing cells (Fig. 6B, C). Notably, all LASVpp underwent type II fusion, similar to LASVpp fusion with endosomes. Thus, irrespective of hLAMP1 expression and the virus entry sites, LASV GPC mediates type II fusion that proceeds through permeabilization of the viral membrane prior to fusion pore opening. Similar to endosomal entry, we observed both instantaneous and delayed release of mCherry following YFP-dequenching (Fig. 6B, C), as well as YFP dequenching without mCherry release (Fig. 6D) deemed as stalled fusion events. mCherry loss from LASVpp was not a result of virus lysis, since these events should not be associated with recovery of YFP signal at low external pH. It should be stressed, however, that, although LASVpp fusion is triggered at the plasma membrane and endosome acidification is blocked by BafA1, pseudoviruses exposed to acidic pH on the cell surface can still be internalized and fuse with neutral endosomes. In fact, the trajectories of some particles traveling several microns, sometimes in directional manner and with velocity reaching 0.8 µm/sec (Fig. 6B-D), are more consistent with endosomal transport of viruses prior to fusion than with viruses stuck at the cell surface.

**Figure 6.**
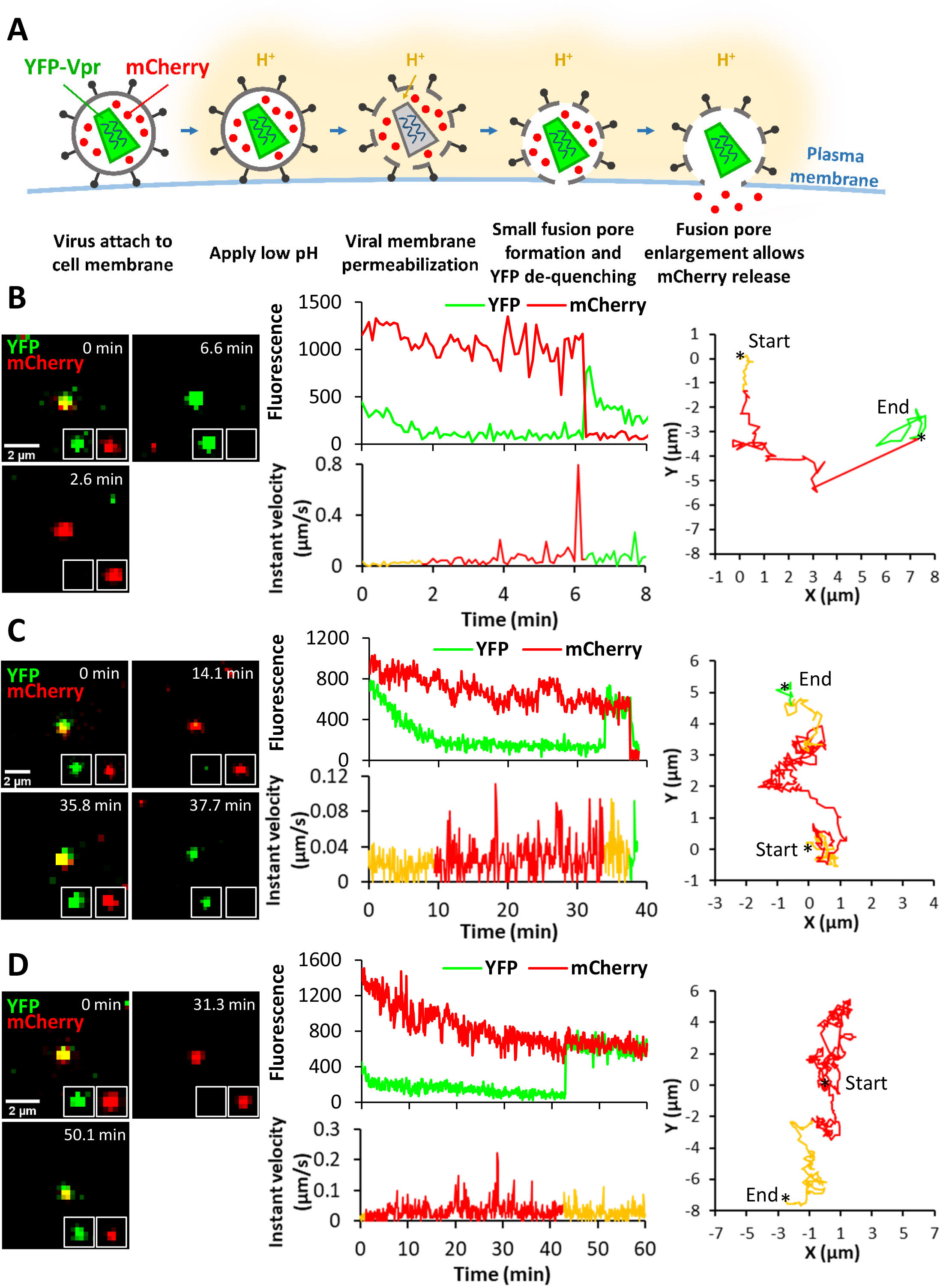
Low pH-forced fusion of single LASVpp with the DF-1 cell plasma membrane. **(A)** Illustration of single mCherry-2xCL-YFP-Vpr labeled LASVpp fusion with the plasma membrane of DF-1 cell induced by exposure to low pH. Viral membrane permeabilization in acidic buffer leads to virus interior acidification and quenching of YFP signal. Subsequent formation of a small fusion pore with the plasma membrane results in virus interior re-neutralization and YFP dequenching. Fusion pore enlargement leads to mCherry release into the cytoplasm. **(B)** Low pH-forced single LASVpp fusion with instant mCherry release. Time lapse images (left), fluorescence traces (middle top), instant velocity (middle bottom) and trajectory (right) of single LASVpp forced fusion in DF-1-LAMP1-d384 cell showing YFP de-quenching and mCherry loss at 6.3 min. (see Movie S4). **(C)** Low pH-forced single LASVpp fusion with delayed mCherry release relative to YFP dequenching. Time lapse images (left), fluorescence traces (middle top), instant velocity (middle bottom) and trajectory (right) of single LASVpp forced fusion with DF-1-pQCXIP cell resulting in YFP dequenching at 34 min and a subsequent loss of mCherry at 37.6 min. (see Movie S5). **(D)** Low pH-forced single LASVpp fusion without mCherry release. Time lapse images (left), fluorescence traces (middle top), instant velocity (middle bottom) and trajectory (right) of single LASVpp fusion with a DF-1-pQCXIP cell showing YFP dequenching at 42.9 min without mCherry loss. (see Movie S6).

Remarkably, forced LASVpp fusion with control DF-1 cells was approximately an order of magnitude more efficient than fusion through an endosomal route (compare Figs. 7A and 5A). The lower efficiency of endosomal fusion might be due to a limited fraction of cell-bound LASVpp being internalized and delivered into acidic endosomes (Fig. S5A), whereas all surface exposed pseudoviruses are synchronously triggered through the forced fusion protocol. It thus appears that endosomal entry of LASVpp into DF-1 cells is limited by slow/inefficient virus uptake. In sharp contrast to a dramatic increase in forced fusion measured by the BlaM assay (Fig. 2C), hLAMP1 expression did not strongly enhance the already efficient forced fusion of single LASVpp (Fig. 7A). Importantly, fusion events associated with simultaneous YFP dequenching and mCherry release were strongly enhanced by hLAMP1 expression as compared to other types of single LASVpp fusion events (Fig. 7A, *Inset*). Thus, hLAMP1 promotes the initial dilation of nascent LASVpp fusion pores triggered at the cell surface to sizes that allow mCherry release. The overall fraction of single LASVpp fusion events associated with mCherry release (delayed or instantaneous) reached ∼25% of all cell-bound particles for DF-1 cells expressing LAMP1-d384 (Fig. 7A).

**Figure 7.**
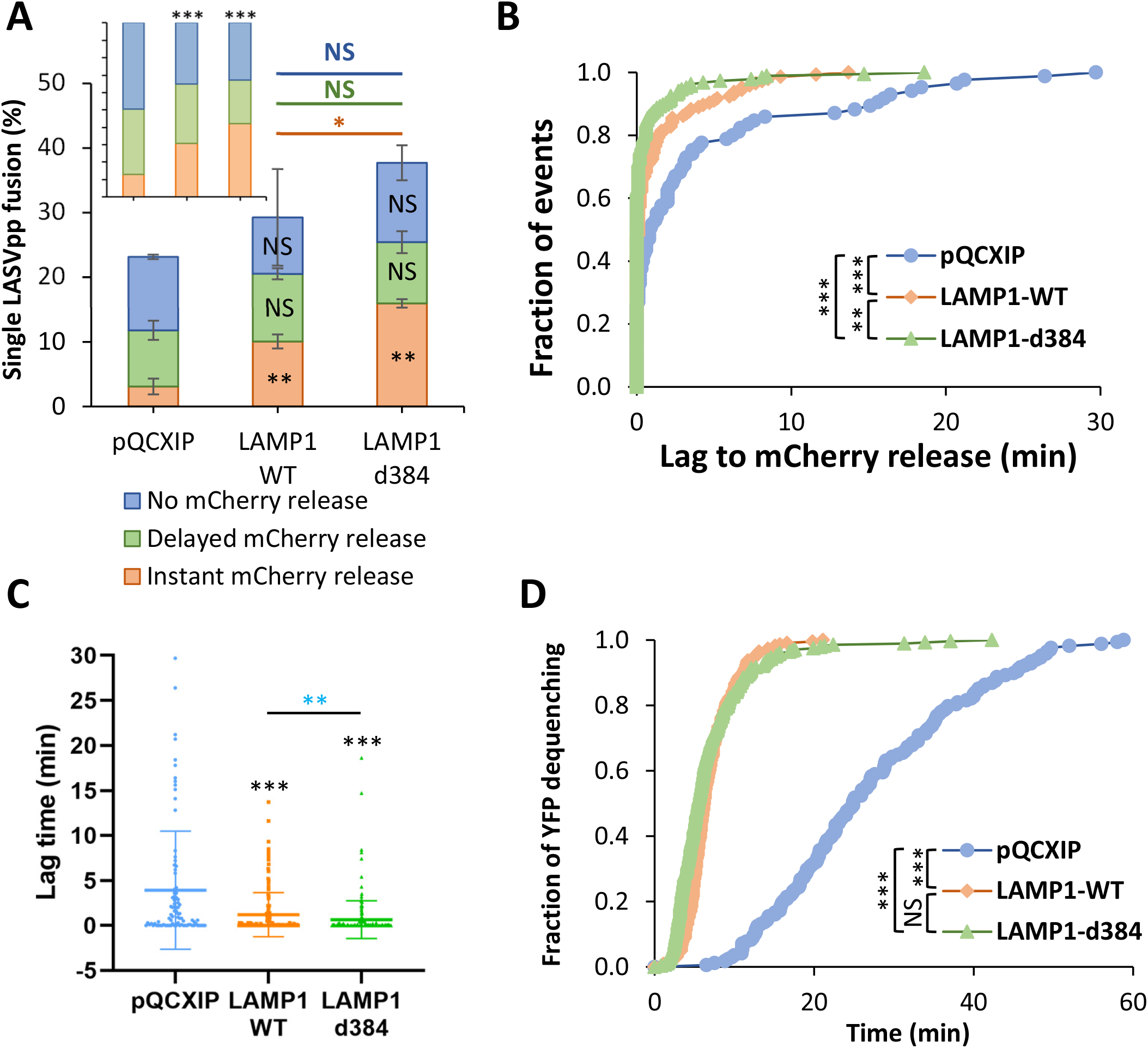
Human LAMP1 enhances low pH-forced fusion of LASVpp with DF-1 cells and dramatically increases the fusion kinetics. **(A)** Efficiencies of low pH-forced single LASVpp fusion events with instant mCherry release, delayed mCherry release and without mCherry release (fusion pore dilation) with DF-1 pQCXIP, LAMP1-WT and LAMP1-d384 cells. Data shown are means ± SD of 2 independent experiments. Black asterisks inside the bars represent significance relative to the vector control. The red, green and blue asterisks show significance levels for the difference between LAMP1-WT and LAMP1-d384 for fusion events with instant mCherry release, delayed mCherry release and without mCherry release respectively. At least 500 cell-bound particles were analyzed for each condition. *Inset*: normalized fractions of each category of fusion. **(B)** The distribution of lag times between small fusion pore formation (YFP dequenching) and pore enlargement (loss of mCherry) for LASVpp fusion with DF-1 pQCXIP, LAMP1-WT and LAMP1-d384 cells. **(C)** Same as in B but plotted as a dot-plot (lines represents means and whiskers are standard deviations). **(D)** The kinetics of small pore formation (YFP dequenching) between single LASVpp and DF-1 pQCXIP, LAMP1-WT and LAMP1-d384 cells using a forced fusion protocol. Data in (A) and (C) are means ± SD of two independent experiments. Blue asterisks show the significance levels for the difference between LAMP1-WT and LAMP1-d384, black asterisks on the top of bars represent significance relative to the vector control. Normalized fractions of different single virus fusion events were analyzed by Fisher’s exact test using R Project. Data of lag time between YFP dequenching and mCherry release was analyzed by non-parametric Mann-Whitney test using GraphPad. Other results were analyzed by Student’s t-test. *, p<0.05; **, p<0.01; ***, p<0.001; NS, not significant.

To further test if hLAMP1 expression promotes fusion pore dilation, we analyzed the lag times between YFP dequenching and mCherry release as a metric for initial enlargement of nascent pores. hLAMP1 expression caused a highly significant decrease in the lag time between YFP-dequenching and mCherry release (Fig. 7B). The average lag time was reduced from 3.9±0.7 min in control cells to 1.2±0.2 and 0.6±0.2 min in WT and mutant hLAMP1 expressing cells, respectively (Fig. 7C), in excellent agreement with the greater fraction of “instantly” dilating pores (Fig. 7A, *Inset*).

As expected for forced fusion that bypasses the need for virus uptake and delivery into acidic endosomes, the kinetics of forced fusion with DF-1 cells was considerably faster than the kinetics of endosomal fusion (Figs. 5B and 7D). While the effect of hLAMP1 expression on the extent of forced fusion was modest, the forced fusion kinetics with hLAMP1 expressing cells was markedly (∼5-fold) faster than with control cells (Fig. 7D). The faster kinetics of fusion and significant reduction in the lag time between YFP dequenching and mCherry loss show that hLAMP1 accelerates forced LASVpp fusion and promotes the initial enlargement of nascent pores. The hLAMP1 effect on the initial pore enlargement in these experiments is in contrast to a non-significant effect on enlargement of LASVpp pores formed in endosomes (Fig. 5A, D). Note, however, that the fraction of “instantly” dilating pores was twice as high upon endosomal fusion relative to the forced fusion (*insets* to Figs. 5A and 7A), indicating a more efficient enlargement of LASVpp fusion pores formed in endosomes.

### The transmembrane domain of hLAMP1 may be required for efficient LASVpp fusion pore dilation

To determine if hLAMP1 must be anchored to a cell membrane to facilitate LASVpp fusion, we expressed and purified the human LAMP1 ectodomain (referred to as soluble LAMP1, sLAMP1) in Expi293F cells (Fig. S6) and assessed its effect on forced LASVpp fusion with DF-1 cells, using BSA as a negative control. Forced LASV fusion in the presence of sLAMP1 primarily exhibited delayed release of mCherry after YFP dequenching or no mCherry release (stalled fusion) (Fig. 8A-D), whereas “instant” pore dilation events were relatively rare (not shown). mCherry release was abrogated in the presence of the fusion inhibitor ST-193 [22] (Fig. S7), demonstrating that these were *bona fide* viral fusion events. Soluble LAMP1 did not significantly increase the overall extent of LASVpp fusion (Fig. 9A), but the fraction of “instant” mCherry release events was slightly but significantly increased (Fig. 9A, *Inset*). A shorter lag between YFP dequenching and mCherry release (Fig. 9B) further supports the notion that sLAMP1 can promote the initial pore dilation, albeit after a measurable delay (Fig. 9A). The average lag time between YFP dequenching and mCherry release decreased from 6.7±1.0 min in control cells to 2.3±0.7 min in the presence of sLAMP1. The lower relative weight of “instantly” dilating fusion pores and the larger fraction of stalled fusion events (∼30% *vs* ∼60%) for forced fusion with sLAMP1-treated DF-1 cells compared to the full-length hLAMP1 expressing cells (Fig. 7A vs Fig. 9A, *Insets*) indicates that the hLAMP1 transmembrane domain may be required for efficient pore enlargement.

**Figure 8.**
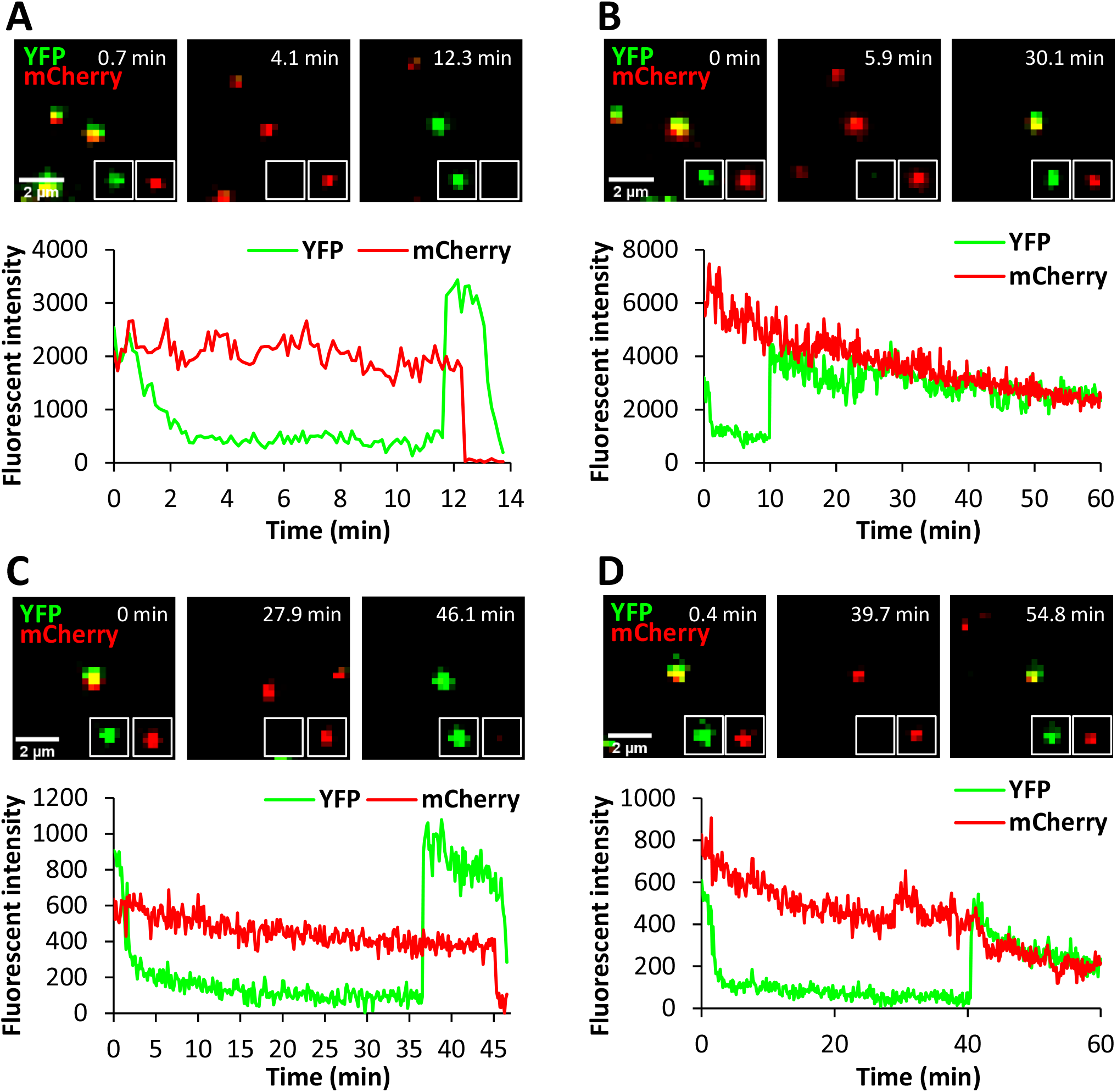
Low pH-forced single LASVpp fusion with DF-1 cells in the presence of soluble LAMP1. LASVpp were bound to cells in the cold, in the presence of 200 µg/ml soluble LAMP1 (sLAMP1) or BSA (control). Single LASVpp fusion with the plasma membrane was initiated by addition of 2 ml of warm pH 5.0 citrate buffer supplemented with 200 µg/ml sLAMP1 or BSA. **(A)** Time lapse images (top) and fluorescence traces (bottom) for single LASVpp fusion with DF-1 cell in the presence of sLAMP1 showing YFP dequenching at 11.7 min and mCherry loss at 12.7 min. **(B)** Time lapse images (top) and fluorescence traces (bottom) of single LASVpp fusion with DF-1 cell in the presence of sLAMP1 showing YFP dequenching at 11.7 min without mCherry loss. **(C)** Time lapse images (top) and fluorescence traces (bottom) of single LASVpp fusion with DF-1 cell in the presence of BSA (control) showing YFP dequenching at 36.4 min and mCherry loss at 46 min. **(D)** Same conditions as in (A), but small fusion pore formation (YFP dequenching) at 40.1 min does not culminate in mCherry release.

**Figure 9.**
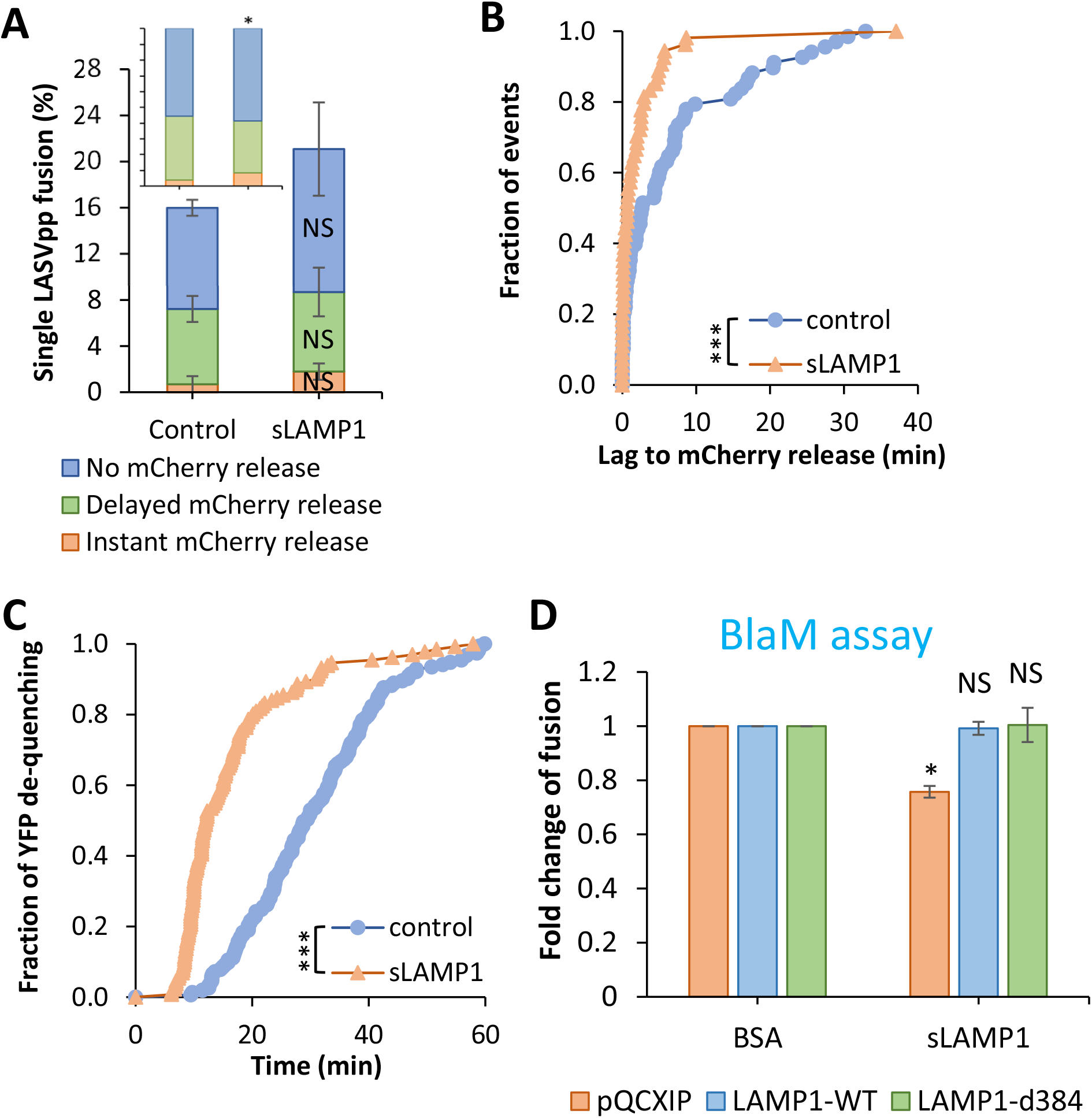
Soluble LAMP1 accelerates low pH-forced single LASVpp fusion with DF-1 cells. **(A)** Efficiencies of low pH-forced single LASVpp fusion events with instant mCherry release, delayed mCherry release and without mCherry release with DF-1 cells in the absence or presence of sLAMP1. Data shown are means ± SD of two independent experiments. *Inset*: normalized fractions of each category of fusion. Asterisk shows significant change relative to the BSA control. **(B)** The distribution of lag times between small fusion pore formation (YFP dequenching) and fusion (loss of mCherry) for forced LASVpp fusion with DF-1 cells in the absence or presence of sLAMP1. **(C)** Kinetics of forced single LASVpp fusion pore formation (YFP dequenching) with DF-1 cells in the absence or presence of sLAMP1. **(D)** Low pH-forced fusion of LASVpp with DF-1 cells in the presence of sLAMP1 measured by the BlaM assay. Cells were pretreated with 0.2 µM BafA1 for 1 h prior to binding pseudoviruses in the cold. Fusion was triggered by applying pH 5.0 citrate buffer at 37 °C for 20 min followed by additional incubation at a neutral pH, 37 °C for 30 min. sLAMP1 (200 µg/ml) or BSA were included throughout virus spinoculation onto cells and low pH triggering of fusion. Data shown are means ± SD of two independent experiments. Normalized fractions of different single virus fusion events were analyzed by Fisher’s exact test using R Project. Data of lag time between YFP dequenching and mCherry release were analyzed by non-parametric Mann-Whitney test using GraphPad. Other results were analyzed by Student’s t-test. *, p<0.05; ***, p<0.001; NS, not significant.

Similar to the effect of the full-length hLAMP1 expression, the kinetics of forced fusion was markedly accelerated by sLAMP1 compared to the BSA control (Fig. 9C). However, the kinetics of forced fusion in the presence of sLAMP1 was more than 2-fold slower than the kinetics of forced fusion with hLAMP1-expressing DF-1 cells (Figs. 7D and 9C). We surmise that the kinetics of forced fusion in the presence of soluble receptor may be limited by the rate of sLAMP1 binding to GPC at low pH.

The marginal effect of sLAMP1 on the initial dilation of forced fusion pores at the cell surface prompted us to test whether the soluble receptor promotes bulk LASVpp fusion, using the BlaM assay. Pseudoviruses were bound to DF-1 cells in the cold, and their fusion with the plasma membrane was triggered by exposing to pH 5.0 in the presence of sLAMP1. Surprisingly, soluble LAMP1 modestly reduced the bulk fusion efficiency with mock-transduced DF-1 cells and had no effect on forced fusion with DF-1 cells expressing LAMP1-WT or LAMP1-d384 (Fig. 9D). This finding implies that, while sLAMP1 accelerates the formation of nascent GPC-mediated fusion pores and marginally increases the probability of initial pore dilation, it fails to promote the formation of fully enlarged fusion pores that can permit the release of the BlaM-Vpr labeled viral cores into the cytoplasm.

## Discussion

Here, we examined the role of LAMP1 in LASV fusion with human and avian cells. Single-virus tracking in live cells combined with functional assays monitoring productive virus fusion/infection provide unprecedented insights into the formation and dilation of nascent fusion pores to functionally relevant sizes. We have previously shown that single LASVpp undergoes type II fusion with endosomes of A549 cells, which entails viral membrane permeabilization (YFP-Vpr signal quenching in acidic endosomes) prior to fusion (YFP dequenching and release of the mCherry content marker) [20]. Type II fusion appears unique to LASVpp GPC, since HIV-1 particles pseudotyped with other viral fusion proteins undergo type I fusion without viral membrane permeabilization prior to fusion [23–30]. Here, we show that LASVpp undergoes type II fusion independent of a target cell (A549 and DF-1 cells) and irrespective of whether fusion occurred in endosomes or at the plasma membrane. The mechanism by which LASV GPC increases the permeability of the viral membrane upon exposure to low pH is currently under investigation.

Unexpectedly, ectopic expression of hLAMP1 has a relatively modest effect on LASV fusion with endosomes across three platforms – LASVpp, LASV-VLP and recombinant LCMV/LASV-GPC viruses, even in avian cells expressing a LAMP1 ortholog that does not support LASV fusion. These results indicate that LASVpp fusion with endosomes is rather efficient, perhaps due to the presence of additional endosomal host factors. Indeed, we have recently shown that the late endosome-resident lipid, bis(monoacylglycero)phosphate (BMP), greatly promotes the dilation of LASV GPC-mediated fusion pores [12]. A modest effect of hLAMP1 expression on LASV fusion and infection could also be due to a limited virus access to this ectopically expressed endosomal receptor due to slow/inefficient virus uptake and delivery into acidic endosomes (Fig. S5A).

In sharp contrast to a modest effect of hLAMP1 expression on LASV entry through an endosomal pathway, we observed a dramatic enhancement of bulk LASV fusion/infection by ectopically expressed hLAMP1 upon forcing fusion at the surface of A549 and DF-1 cells (Fig. 2 and Fig. S2). This striking difference in hLAMP1-dependence of fusion between the conventional and forced LASV entry pathways is suggestive of key differences in the requirements for productive fusion with the plasma membrane *vs* endosomes. Clearly, hLAMP1 plays a critical role in LASV fusion with the plasma membrane which likely lacks requisite endosomal co-factors that augment LASV fusion. Interestingly, forced LCMV-LASV GPC infection was much less dependent on hLAMP1 expression than forced LASVpp infection (Figs. 2E and 3C). This difference may reflect a more efficient dilation of fusion pores in the context of a recombinant LCMV/LASV virus, perhaps because of a greater surface density of GPCs, as compared to HIV-based pseudoviruses.

Inefficient LASVpp pore enlargement at the cell surface is further supported by the strikingly different effects of hLAMP1 expression on the forced fusion and infection, as measured by the BlaM and luciferase assays in DF-1 cells, as compared to a modest increase of forced single virus fusion (Fig. 2C, E *vs* Fig. 7A). This differential effect is likely related to distinct requirements for the functional pore size for the bulk BlaM/infectivity assays compared to a single-virus fusion assay. It should be stressed that the latter assay cannot monitor fusion pore dilation beyond sizes that allow mCherry release; once mCherry is released, there is no way of knowing whether a fusion pore further expanded, remained relatively small or collapsed. For this reason, single virus imaging reports the formation and initial dilation of nascent fusion pores and not the formation of fully enlarged pores that allow for productive infection. While single virus tracking enables highly sensitive detection of nascent fusion pores in a sub-nanometer/nanometer range, the BlaM signal and productive infection likely require full dilation of pores to release the HIV-1 capsid core (∼100 nm [31]) into the cytoplasm. It should be noted that a previous study [32] has detected inadvertent cleavage of a BlaM-Vpr construct incorporated into HIV-1 virions by the viral protease and concluded that free BlaM molecules (which can diffuse through a small fusion pore) are responsible for cleavage of a BlaM substrate in infected cells. However, due to the lack of detectable free BlaM in our virus samples (Fig. S1A), it is more likely that BlaM signal originates from the HIV-1 core-incorporated BlaM-Vpr, which must be released into the cytoplasm through a fully enlarged fusion pore. This notion is further supported by a marked hLAMP1-mediated enhancement of the BlaM signal upon forced fusion of LASV-VLPs containing NP-BlaM chimera, which is not cleaved inside VLPs (Fig. S1B). This result suggests that the hLAMP1 effect on dilation of LASV GPC-mediated fusion pores is independent of virus context (HIV-1 core *vs* arenavirus VLP).

Besides the effect of hLAMP1 on fusion pore dilation, overexpression of this intracellular LASV receptor strongly accelerates GPC-mediated fusion with endosomes and forced fusion at the cell surface. The faster kinetics of LASVpp fusion with endosomes may be due to a less acidic pH threshold for fusion with LAMP1 expressing membranes [10, 12] which enables virus entry from earlier endosomal compartments. It appears that GPC-hLAMP1 interactions at low pH can trigger more concerted conformational changes in the LASV fusion protein, leading to a more efficient and quick formation of functional prefusion complexes, likely consisting of several activated trimers. The fusion-accelerating effect of hLAMP1 appears independent from the transmembrane domain of LAMP1, since soluble LAMP1 also accelerates forced LASVpp fusion at the cell surface, whereas the transmembrane domain may be required for efficient pore dilation (Fig. 9D). Thus, the hLAMP1-mediated acceleration of formation of nascent fusion pores may be distinct from its promotion of LASV pore dilation. It is currently unclear whether the transmembrane domain of hLAMP1 has a specific role in enlargement of LASV GPC-mediated fusion pores, or it simply anchors the ectodomain to a target membrane, thereby increasing the local surface density of this receptor for more efficient triggering of GPC refolding at low pH.

To reconcile the above observations, we propose the following model for LASV fusion. LASVpp are capable of forming small pores with endosomes of less permissive DF-1 cells and with the plasma membrane upon low pH exposure in the absence of hLAMP1. However, the majority of LASVpp fusion pores, especially those formed through the forced fusion protocol, does not enlarge to sizes that allow the cytosolic delivery of BlaM-Vpr or infection. In fact, stalled fusion pores (YFP dequenching without mCherry loss) formed upon forcing virus fusion by low pH represent nearly half of events for control DF-1 cells (Fig. 7A), supporting the notion of incomplete dilation of at least a fraction of fusion pores. Based on the dramatic increase in forced fusion and infection upon ectopic hLAMP1 expression (Fig. 2), this receptor profoundly increases the probability of formation of functional fusion pores with suboptimal targets, such as the plasma membrane. The much-accelerated LASVpp fusion kinetics observed in our experiments and a shift in the pH-optimum for fusion toward less acidic pH [10, 12] in the presence of hLAMP1 imply that this intracellular receptor favors the initiation of LASV fusion with earlier endosomal compartments. It appears, however, that productive LASV fusion with endosomes is augmented by additional host factors, such as the BMP, that promote dilation of LASV fusion pores [12]. Considering the difficulties associated with controlling the conditions (e.g., pH) in endosomes and dissecting the roles of endosomal co-factors in viral fusion, redirecting the viral fusion to a less optimal target membrane, e.g., cell plasma membrane, is a viable strategy to unraveling the effects of host co-factors on productive virus fusion.

## Materials and Methods

### Cell lines and transfection

Human embryonic kidney 293T/17 cells, human lung epithelial A549 cells and chicken embryonic fibroblast DF-1 cells were obtained from ATCC (Manassas, VA, USA). A549 and DF-1 cells stably expressing wild-type and the Ala384-deleted LAMP1 mutant (LAMP1-WT and LAMP1-d384) were generated by transduction with pQCXIP retroviral vectors (Clontech, Mountain View, CA) encoding LAMP1-WT and LAMP1-d384. pQCXIP empty vector was used to generate control A549 and DF-1 vector cell lines. NuExpi293F cells were a gift of Dr. Jens Wrammert (Emory University).

Except for Expi293F cell, cells were maintained in high glucose Dulbecco’s Modified Eagle Medium (DMEM; Mediatech, Masassas, VA, USA) containing 10% heat-inactivated Fetal Bovine Serum (FBS; Atlanta Biologicals, Flowery Branch, GA, USA) and 1% penicillin/streptomycin (GeminiBio, West Sacramento, CA, USA). For HEK 293T/17 cells, the growth medium was supplemented with 0.5 mg/ml G418 (Genesee Scientific, San Diego, CA, USA). Expi293 cells were maintained in Expi293^TM^ Expression Medium (Life Technologies Corporation, NY, USA).

HEK293T/17 cells were transfected with JetPRIME transfection reagent (Polyplus-transfection, Illkirch-Graffenstaden, France) according to the manufacturer’s instructions. Expi293F cells were transfected with Sinofection Transfection Reagent (SinoBiological, Beijing, P.R. China) according to the manufacturer’s instructions.

### Cloning and stable cell lines construction

To generate the plasmids pQCXIP-LAMP1-WT and pQCXIP-LAMP1-d384, LAMP1-WT and LAMP1-d384 segments were amplified from pcDNA-LAMP1-WT and pcDNA-LAMP1-d384 (a gift of Dr. Ron Diskin, Weizmann Institute) respectively by PCR (forward primer: GCACCGGTATGGCGGCCCCCGGCAGCGC; reverse primer for LAMP1 WT: CGCGGATCCCTAGATAGTCTGGTAGCCTGCG; reverse primer for LAMP1-d384: CGCGGATCCCTAGATAGTCTGGTAGCCGTG) and inserted into pQCXIP by *AgeI*/*BamHI* restriction enzyme digestion and ligation.

JUNV-NP and JUNV-Z plasmids were a gift of Dr. Jack H. Nunberg (University of Montana). To generate the plasmid JUNV-NP-BlaM, β-lactamase segments were amplified by PCR (forward primer: CCATGAGGAGTGTTCAACGAAACACAGTTTTCAAGGTGGGAAGCTCCGGCGACCCAGAAACGCTGGTGAAAG; reverse primer: GGTCAGACGCCAACTCCATCAGTTCATCCCTCCCCAGGCCGGAGCTGCCCCAATGCTTAATCAGTGAGGCACC) and inserted into JUNV-NP at NP amino acid residue 93 by QuickChange PCR with the β-lactamase segments served as megaprimers.

To create pQCXIP vector cells and cells stably expressing LAMP1-WT and LAMP1-d384, pseudotyped retroviruses were produced by transfecting HEK293T/17 cells with pQCXIP/pQCXIP-LAMP1-WT/pQCXIP-LAMP1-d384, MLV-Gag-Pol and VSV-G using JetPRIME transfection reagent. Supernatants were harvested 36-48 h post-transfection and filtered with 0.45 µm filter to remove cell debris and virus aggregates. A549 or DF-1 cells were infected with the pseudoviruses and selected in a growth medium containing 1.5 or 2.5 µg/ml puromycin at 24 h after infection for A549 and DF-1 cells, respectively.

### Immunostaining for LAMP1

To confirm the expression of LAMP1, cells were seeded on 8-well chambered coverslips (LabTek, MA, USA). Cells were washed with PBS^++^, fixed with 4% PFA (Electron Microscopy Sciences, PA, USA) at room temperature for 15 min, permeabilized with 125 μg/ml digitonin (Research Products International, IL, USA) at room temperature for 15 min, and incubated with 10 µg/ml of mouse anti-hLAMP1 H4A3 antibody (Abcam Inc, Waltham, MA, USA) at room temperature for 1 hour. To assess the LAMP1 expression level on the cell surface, cells were incubated with anti-hLAMP1 antibody in the cold before PFA fixation, without cell permeabilization. Cells were then washed and incubated with 1 µg/ml AlexaFluor647 Donkey Anti-Mouse IgG (H+L) (Thermo Fisher Scientific Corporation, OR, USA) at room temperature for 45 min. Cell nuclei were stained with 10 µM Hoechst-33342 (Molecular Probes, OR, USA). Images were acquired on a Zeiss LSM 880 confocal microscope using a plan-apochromat 63X/1.4NA oil objective. LAMP1 expression level were quantified by integrating the fluorescence intensity per cell after background subtraction using ImageJ.

### Soluble LAMP1 expression and purification

Expi293F cells were transfected with pHLsec-Lamp1 fragment, a kind gift of Juha T. Huiskonen (University of Oxford, Oxford, UK), followed by three days incubation at 37 °C in the presence of 1 µg/ml of kifunensine (R&D Systems, MN, USA). Supernatant was collected and combined with half volumes of the supernatant of binding buffer (25 mM HEPES, 150 mM NaCl, pH 7.2). Soluble LAMP1 fragment (sLAMP1) was purified by Ni-NTA affinity chromatography. The elution was desalted and diluted in PBS and concentrated to 5 mg/ml. Purity of sLAMP1 was assesses by SDS-PAGE and Coomassie Blue staining and Western-blotting.

### Pseudovirus and VLP production

HEK293T/17 cells were seeded in growth medium supplemented with 10% FBS a day before transfection. For the bulk virus-cell fusion assay, LASVpp carrying β-lactamase-Vpr chimera (BlaM-Vpr) were produced by transfecting HEK293T/17 cells with Lassa GPC, pR9ΔEnv, BlaM-Vpr and pcRev plasmids. LASV-VLPs carrying β-lactamase-NP (JUNV candid-1) chimera (JUNV-BlaM-NP) were produced by transfecting HEK293T/17 cells with Lassa GPC, JUNV-NP, JUNV-BlaM-NP and JUNV-Z plasmids. For single virus fusion experiments, dual-labeled LASVpp were produced by transfecting HEK293T/17 cells with Lassa GPC, pR9ΔEnv, mCherry-2xCL-YFP-Vpr [19] and pcRev plasmids. Supernatants were collected at 36-48 h post-transfection, filtered with 0.45 µm filter to remove cell debris and virus aggregates. LASVpp-BlaM virus was concentrated 10 times with Lenti-X concentrator (Clontech Laboratories, Mountain View, CA, USA) according to the manufacturer’s instructions. The β-lactamase in LASVpp-BlaM and LASV-VLP-BlaM incorporation and cleavage were examined by Western blotting using anti-β-lactamase antibody (QED Bioscience Inc, CA, USA).

### Virus-cell fusion assay

Target cells were seeded in phenol red-free DMEM supplemented with 10% FBS. LASVpp-BlaM or LASV-VLP-BlaM particles were bound to cells by centrifugation at 4 °C for 30 min at 1550xg. Cells were washed with cold phenol red-free DMEM/10% FBS buffered by 20 mM HEPES (GE Healthcare Life Sciences) to remove unbound viruses. Viral fusion was initiated by shifting to 37 °C for 2 h, after which time, cells were placed on ice, loaded with the CCF4-AM substrate (Life Technologies), and incubated overnight at 11 °C. In control experiments, fusion was performed in the presence of 40 mM NH_4_Cl to raise endosomal pH and block virus entry. The cytoplasmic BlaM activity (ratio of blue to green fluorescence) was measured using a SpectraMaxi3 fluorescence plate reader (Molecular Devices, Sunnyvale, CA, USA). The background BlaM signal (blue/green ratio) in the presence of NH_4_Cl was subtracted from the fusion signal before calculating the fold-increase in fusion upon hLAMP1 expression. Note that, although the LASVpp infection of DF-1 cells was markedly lower than of A549 cells, DF-1 cells produced more robust BlaM signals for the same MOI. This is likely caused by better loading and/or retention of the CCF4-AM substrate by DF-1 cells.

For low pH-forced fusion at the plasma membrane, cells were pre-treated with 0.2 µM Bafilomycin A1 for 1 h. LASVpp-BlaM or LASV-VLP-BlaM particles were bound to A549 and DF-1 cells in the cold. Viral fusion was triggered by incubating the cells with citrate pH 5.0 buffer (50 mM citrate buffer, 5 mM KCl, 2 mM CaCl_2_, 90 mM NaCl, pH 5.0, 300 mOsM) for 20 min and further incubated in phenol red-free DMEM/10% FBS for 30 min at 37 °C. To assess the effect of sLAMP1 on forced LASVpp fusion, 200 µg/ml of sLAMP1 or BSA (control) was included throughout virus spinoculation onto cells and low-pH triggering of fusion. The resulting BlaM activity was measured, as described above for the conventional viral entry protocol.

To avoid saturation of the BlaM signal and to make sure it is in the lineage range, different dilutions of virus stock (MOIs) were used under different conditions. For LASVpp-BlaM fusion through endosomal pathway, MOI of 0.1 and 0.05 were used for A549 and DF-1 cell lines, respectively. For the LASVpp-BlaM forced fusion, we used MOI of 5, 0.5 and 0.05 for A549-pQCXIP, DF-1-pQCXIP and for A549 or DF-1 cells expressing LAMP1-WT or LAMP1-d384, respectively. Since the titer for LASV-VLP-BlaM particles could not be determined, we used the following dilutions of the virus stock to infect different cell lines. For LASV-VLP-BlaM entry through endosomal fusion, 2x and 3x less VLPs were used to infect hLAMP1-expressing cells than control A549 and DF-1 cells, respectively. For LASV-VLP-BlaM entry through a forced pathway, 3x and 50x less VLP was used to infect LAMP1 expressing cells than control A549 and DF-1 cells, respectively.

### Infectivity assay for LCMV/LASV-GPC recombinant virus

Target cells were seeded in DMEM supplemented with 10% FBS and grown to 70% confluency. LCMV/LASV-GPC viruses (MOI 0.01) were bound to cells by centrifugation at 4 °C for 30 min at 1550xg. Cells were washed with cold phenol red-free DMEM/10% FBS buffered by 20 mM HEPES (GE Healthcare Life Sciences) to remove unbound viruses. Infection was allowed by incubating at 37 °C for 20 h. For the infection through forced pathway, cells were pretreated with 0.2 µM Bafilomycin A1 for 1 hour. LCMV/LASV-GPC virus (MOI 0.1) were bound to cell in the cold. Viral fusion was triggered by incubating the cells with citrate pH 5.0 buffer for 20 min at 37 °C and further cultured in phenol red-free DMEM/10% FBS for 20 h at 37 °C. Cells were washed with PBS++, fixed with 4% PFA at room temperature for 15 min, permeabilized with 0.3% TritonX-100 (Sigma, St. Louis, MO, USA) at room temperature for 15 min, and incubated with 5 µg/ml rat anti-LCMV NP antibody (Bio X Cell, Lebanon, NH, USA) at room temperature for 1 hour. Cells were then washed and incubated with 1 µg/ml Donkey anti-Rat polyclonal Dylight 488-conjugated antibody (Thermo Fisher Scientific Corporation, OR, USA) at room temperature for 45 min. Cell nuclei were stained with 10 µM Hoechst-33342 (Molecular Probes, OR, USA). Images were acquired by Cytation 3 imaging plate reader (Agilent Technologies, Santa Clara, CA, USA), cell nuclei and LCMV NP expression in infected cells were detected by Gen5 software (Agilent Technologies, Santa Clara, CA, USA). Infection was calculated as the fraction of infected cell.

### LASV pseudovirus infectivity assay

Target cell were seeded in phenol red-free DMEM supplemented with 10% FBS. Luciferase-encoding LASVpp were bound to cells by centrifugation at 4 °C for 30 min at 1550xg (MOI=0.1 for A549 cells and MOI=10 for DF-1 cells). Unbound virus was removed by washing with cold medium and the cells were incubated with 50 µl growth medium at 37 °C to allow infection. For the infection through forced pathway, cells were pretreated with 0.2 µM Bafilomycin A1 for 1 h. LASVpp were bound to cell in the cold (MOI=1 for A549 cells and MOI=10 for DF-1 cells). Viral fusion was triggered by incubating the cells with citrate pH 5.0 buffer for 20 min followed by further incubation with 50 µl growth medium at 37 °C to allow infection. Thirty-six hours post-infection, cells were incubated with 50 µl Bright-Glo^TM^ luciferase substrate (Promega, Madison, WI) at room temperature for 5 min, the luciferase activity was measured by a TopCount NXT plate reader (PerkinElmer Life Sciences, Waltham, MA, USA).

### Single virus imaging in live cell

Target cells were seeded in 35 mm collagen coated glass-bottom Petri dishes (MatTek, MA, USA) in Fluorobrite DMEM (Life Technologies Corporation, NY, USA) containing 10% FBS, penicillin, streptomycin and L-glutamine 2 days before imaging. The binding of dual-labeled LASVpp to cells (MOI of 0.1) was facilitated by centrifugation at 4 °C, and cells were washed with cold PBS^++^ to remove unbound virus. Virus entry and fusion were initiated by adding 2 ml of pre-warmed Fluorobrite DMEM containing 10% FBS immediately prior to imaging cells on a DeltaVision microscope equipped with a temperature, humidity and CO_2_-controlled chamber for 2 h. Every 6 sec, 4 Z-stacks spaced by 1.5 μm were acquired to cover the thickness of cells using Olympus 60x UPlanFluo /1.3 NA oil objective (Olympus, Japan). For low pH-forced fusion at the plasma membrane, cells were pre-treated with 0.2 µM Bafilomycin A1 (Sigma, St. Louis, MO, USA) for 1 h. Forced fusion was initiated by adding 2 mL citrate pH 5.0 buffer with 0.2 µM Bafilomycin A1, followed by imaging for 1 h at 37 °C. To assess the effect of sLAMP1 on LASVpp forced fusion, 200 µg/ml of sLAMP1 or BSA (control) was added to virus during spinoculation onto cells and included in a citrate pH 5.0 buffer during virus entry and imaging. In control experiments, 10 µM of ST-193 (MedChemExpress, NJ, USA) was applied to the mixture virus during spinoculation onto cells and the inhibitor and 0.2 mg/ml of sLAMP1 were included in citrate pH 5.0 buffer and maintained throughout imaging.

The acquired time-lapse Z-stack images were converted to maximum intensity projections for single particle tracking. Single fusion events were annotated using the ImageJ ROI manager tool. The times of YFP dequenching and mCherry loss, which occurred in a single image frame, were determined visually as the time of color change. Representative single virus fusion events were tracked using ICY image analysis software (icy.bioimageanalysis.org). The labeled pseudoviruses were identified by Spot Detection plugin and tracked using Spot Tracking plugin to determine the fluorescence intensity over time, particle trajectory and instant velocity.

### Statistical analysis

Data of lag time between YFP dequenching and mCherry release was analyzed by non-parametric Mann-Whitney test using GraphPad. Normalized fractions of different single virus fusion events were analyzed by Fisher’s exact test using R Project. Other results were analyzed by Student’s t-test using Excel. Unless stated otherwise, *, p<0.05; **, p<0.01; ***, p<0.001; NS, not significant.

## Acknowledgements

The authors wish to thank Jack Nunberg (University of Montana) for the gift of JUNV-NP and JUNV-Z plasmids and cloning advice, Ron Diskin (Weizmann Institute) for WT and mutant human LAMP1 expression vectors, and Juha Huiskonen (University of Oxford) for the gift of sLAMP1 expression vector. We are also grateful to Xiangyang Guo for advice on sLAMP1 expression and purification, Baek Kim (Emory University) for access to Cytation 3 plate reader, and the members of Melikyan lab for critical reading of the manuscript and helpful comments. This work was supported by the NIH R01 AI053668 grant to G.B.M. The funders had no role in study design, data collection and analysis, decision to publish, or preparation of the manuscript

## Supplementary Figure Legends

**Figure S1. BlaM-Vpr or NP-BlaM are not cleaved in pseudoviruses (LASVpp-BlaM) and virus-like particles (LASV-VLP-BlaM).** BlaM-Vpr **(A)** or NP-BlaM **(B)** in LASVpp-BlaM and LASV-VLP-BlaM particles, respectively, were examined by Western blotting using anti-β-lactamase antibody. LASVpp carrying mCherry-YFP-Vpr and LASV-VLP-NP were used as negative controls for non-specific with signal.

**Figure S2. LAMP1 expression enhances LASV-VLP fusion with A549 and DF-1 cells. (A)** LASV-VLP-BlaM fusion with A549 and DF-1 cells. LASV-VLP entry through an endosomal pathway was initiated by pre-binding the VLP in the cold, shifting to 37 °C and incubating for 2 h. **(B)** Low pH-forced fusion of LASV-VLP with A549 and DF-1 cells. Cells were pretreated with 0.2 µM BafA1 for 1 h prior to binding the VLPs in the cold. Fusion was triggered by applying pH 5.0 citrate buffer at 37 °C for 20 min followed by additional incubation in a neutral pH medium at 37 °C for 30 min. Data shown are means ± SD of three independent experiments. Data were analyzed by Student’s t-test. **, p<0.01; ***, p<0.001.

**Figure S3. Fusion of single LASVpp with the DF-1 cell**. **(A)** Time lapse images (top) and fluorescence traces (bottom) of single LASVpp fusion with a DF-1-pQCXIP cell showing YFP quenching at 29.5 min and YFP dequenching/mCherry loss at 46.5 min corresponding to virus interior acidification and fusion, respectively. **(B)** Time lapse images (top) and fluorescence traces (bottom) of single LASVpp fusion with a DF-1-LAMP1-d384 cell showing YFP quenching at 5 min and YFP dequenching/mCherry loss at 25 min, indicating virus interior acidification and fusion, respectively.

**Figure S4. Saponin lysis of LASVpp labeled with mCherry-2xCL-YFP-Vpr attached to a coverslip.** Images (left panel) and quantification (right panel) of virus lysis (loss of mCherry from YFP-Vpr labeled particles) before and 10 min after application of saponin. A small fraction (∼15%) of immature HIV-1 pseudoviruses retained mCherry. Data are means ± SD from 4 image fields analyzed.

**Figure S5. LAMP1 expression doesn’t affect single LASVpp endocytosis and viral membrane permeabilization. (A)** Fraction of single LASVpp exhibiting YFP quenching in DF-1 pQCXIP, LAMP1-WT and LAMP1-d384 cells. Data shown are means ± SD of 5 independent experiments. **(B)** Kinetics of the YFP quenching of single LASVpp in control and hLAMP1 expressing DF-1 cells. Data were analyzed by Student’s t-test. *, p<0.05; **, p<0.01; NS, not significant.

**Figure S6. SD**S-PAGE and Western-blot of purified sLAMP1.

**Figure S7. ST-193 inhibits single LASVpp fusion forced by low-pH.** LASVpp were bound to DF-1 cells in the cold, in the presence of 200 µg/ml soluble LAMP1 with or without 10 µM of ST-193. Single LASVpp fusion with the plasma membrane was initiated by addition of 2 ml of warm pH 5.0 citrate buffer supplemented with 200 µg/ml sLAMP1 with or without 10 µM of ST-193. Graph show efficiencies of low pH-forced single LASVpp fusion events with instant mCherry release, delayed mCherry release and without mCherry release with DF-1 cells in the absence of sLAMP1 with or without ST-193. Data shown result of 1 independent experiment.

## Supplementary Movie Legends

**Movie S1. LASVpp fusion event with instant viral content release following YFP dequenching.** Single LASVpp exhibits YFP quenching (white arrow) at 31.3 min followed by YFP dequenching/mCherry loss at 34.7 min corresponding to virus interior acidification and fusion, respectively. The frame rate is slowed 10-fold around the YFP quenching and dequenching events (29:36–32:12 min and 33:42–35:54 min, respectively) to better illustrate these steps of viral fusion. Movie is related to Fig. 4B.

**Movie S2. LASVpp fusion event with delayed viral content release following YFP dequenching.** Single LASVpp exhibits YFP quenching (white arrow) at 39.7 min followed by YFP dequenching at 42.7 min and mCherry loss at 43.2 min corresponding to virus interior acidification, small fusion pore formation and fusion pore dilation, respectively. The frame rate is slowed 4-fold around the YFP quenching, dequenching and mCherry release events (42:24-46:54). Movie is related to Fig. 4C.

**Movie S3. LASVpp fusion events without viral content release.** Single LASVpp exhibits YFP quenching (white arrow) at 42.6 min followed by YFP dequenching at 47.4 min corresponding to virus interior acidification and small fusion pore formation, respectively. The frame rate is slowed 4-fold around the YFP quenching and dequenching events (41:24-48:24 min). Movie is related to Fig. 4D.

**Movie S4. Low pH-forced LASVpp fusion with instant viral content release.** Single LASVpp exhibits YFP quenching (white arrow) at 1.3 min followed by YFP dequenching/mCherry loss at 6.3 min corresponding to virus interior acidification and fusion, respectively. Movie is related to Fig. 6B.

**Movie S5. Low pH-forced LASVpp fusion with delayed viral content release.** Single LASVpp exhibits YFP quenching (white arrow) at 9.5 min followed by YFP dequenching at 34 min and mCherry loss at 37.6 min corresponding to virus interior acidification, small fusion pore formation and fusion pore dilation, respectively. The frame rate is slowed 10-fold around the YFP quenching and dequenching events (5:54-11:00 min and 32:54-38:24min, respectively). Movie is related to Fig. 6C.

**Movie S6. Low pH-forced LASVpp fusion without viral content release.** Single LASVpp exhibits YFP quenching (white arrow) at 1.4 min followed by YFP dequenching at 42.9 min corresponding to virus interior acidification and small fusion pore formation, respectively. The frame rate is slowed 20-fold around the YFP quenching and dequenching events (0-2:00 min and 42:00-44:00 min, respectively). Movie is related to Fig. 6D.

